# Closed-loop microstimulations of the orbitofrontal cortex during real-life gaze interaction enhance dynamic social attention

**DOI:** 10.1101/2023.12.18.572176

**Authors:** Siqi Fan, Olga Dal Monte, Amrita R. Nair, Nicholas A. Fagan, Steve W. C. Chang

**Affiliations:** Department of Psychology, Yale University, New Haven, CT 06520, USA; The Rockefeller University, New York, NY 10065, USA; Department of Psychology, University of Turin, 10124 Torino, Italy; Department of Neuroscience, Yale University School of Medicine, New Haven, CT 06510, USA; Kavli Institute for Neuroscience, Yale University School of Medicine, New Haven, CT 06510, USA; Wu Tsai Institute, Yale University, New Haven, CT 06510, USA

**Author notes:** Contributed equally (Siqi Fan, Olga Dal Monte).

**Keywords:** Closed-loop microstimulation, dynamic social attention, real-life social gaze interaction, naturalistic social behavior, prefrontal cortex, orbitofrontal cortex, dorsomedial prefrontal cortex, anterior cingulate cortex, non-human primates

## Abstract

The prefrontal cortex is extensively involved in social exchange. During dyadic gaze interaction, multiple prefrontal areas exhibit neuronal encoding of social gaze events and context-specific mutual eye contact, supported by a widespread neural mechanism of social gaze monitoring. To explore causal manipulation of real-life gaze interaction, we applied weak closed-loop microstimulations that were precisely triggered by specific social gaze events to three prefrontal areas in monkeys. Microstimulations of orbitofrontal cortex (OFC), but not dorsomedial prefrontal or anterior cingulate cortex, enhanced momentary dynamic social attention in the spatial dimension by decreasing distance of one’s gaze fixations relative to partner monkey’s eyes. In the temporal dimension, microstimulations of OFC reduced the inter-looking interval for attending to another agent and the latency to reciprocate other’s directed gaze. These findings demonstrate that primate OFC serves as a functionally accessible node in controlling dynamic social attention and suggest its potential for a therapeutic brain interface.

## Introduction

The prefrontal cortex evolved to process a wide range of information in order to adaptively guide behaviors in complex environments ^1^. For social animals, it has been hypothesized that the prefrontal cortex, and other brain regions, prioritize social information to successfully navigate volatile social environments involving multiple conspecifics in group settings ^2-5^. In many primate species, social gaze plays a pivotal role in conveying essential social information ^6^, and several prefrontal brain regions are known to exhibit selective neural activity for social gaze interaction ^7, 8^. While multiple subregions in the primate temporal and posterior parietal cortices, including the gaze-following patch, have been widely implicated in the perceptual aspects of social gaze ^9-12^, the prefrontal subregions are theorized to play critical functions in integrating social, affective, and motivational information to enable appropriate social gaze processing ^13^.

The neural systems involved in social gaze interaction must distinguish social from non-social gaze events, and also mark significant interactive events such as mutual eye contact, in order to regulate social behaviors. This is likely facilitated by the continuous monitoring of one’s own gaze and other’s gaze over time. Recent research in pairs of rhesus macaques has demonstrated that a large proportion of individual neurons in the prefrontal cortex, including the orbitofrontal cortex (OFC), the dorsomedial prefrontal cortex (dmPFC), and the gyrus of anterior cingulate cortex (ACCg), exhibit robust neural representations for gaze fixations directed toward the eyes and the face of a conspecific partner and for context-specific mutual eye contact events ^8^. Importantly, a substantial proportion of cells in these areas were found to parametrically track the Euclidian distance of one’s own gaze fixations in space relative to a partner monkey’s eyes and the distance of the partner’s gaze fixations relative to one’s own eyes ^8^. Dynamic changes in these gaze distance variables provide information on the proximity of gaze fixations of interacting individuals to one another. This information becomes particularly valuable for computing interactive gaze events, such as mutual eye contact or joint attention, when gaze distance variables for self and other converge to specific values. Thus, this parametric representation of gaze-related distances in individual neurons is a noteworthy finding as it can provide a moment-by-moment index of social attention during an ongoing social interaction, possibly serving as a simple, yet elegant, mechanism of social gaze monitoring. This type of gaze-distance coding is not specific to social information processing, however. In OFC, it has been shown that a large proportion of neurons encode the gaze fixation distance from a value-predicting cue on the screen during free viewing ^14^, providing a potentially shared mechanism linking gaze position and reward valuation in both social and non-social contexts.

Nevertheless, a lingering question remains: do neural populations in OFC, dmPFC, or ACCg causally contribute to dynamic social attention? To address this question, here we applied weak, real-time, closed-loop microstimulations unilaterally to each of the three prefrontal areas upon the precise moment when the stimulated monkey fixated on partner monkey’s eyes. Compared to sham stimulations, microstimulations of the OFC facilitated social attention in the spatial dimension by decreasing the average distance of one’s own gaze fixations relative to partner’s eyes. Importantly, this effect was more pronounced for gaze fixations in the contralateral visual field and specific to attending to social stimuli. Moreover, microstimulations of the OFC also exerted an influence in the temporal dimension of social attention by reducing the inter-looking interval for attending to partner’s face as well as reducing the latency to reciprocate partner’s directed gaze. Thus, microstimulations of OFC had a dual impact on both spatial and temporal aspects of dynamic social attention by facilitating focal visual attention around another social agent and promoting reciprocal gaze exchanges. These findings highlight the primate OFC as a causal node in controlling dynamic social attention.

## Results

Two unique pairs of rhesus macaques (M1: stimulated monkey or ‘self’, monkeys L and T; M2: partner monkey or ‘other’, monkey E) engaged in spontaneous face-to-face social gaze interaction ^8, 15^ while the gaze positions of both monkeys were continuously and simultaneously tracked at high temporal and spatial resolution. To examine the causal moment-by-moment contributions of different prefrontal areas in live social gaze interaction, we applied weak, real-time, closed-loop microstimulations (75 µA, 100 Hz, 200 msec; Methods) with a probability of 50% (half microstimulation trials and half sham trials) contingently upon the moment when the stimulated monkey fixated on the partner monkey’s eyes in the live social gaze condition (**Fig. 1a**, left; Methods) or on a random dot motion (RDM) stimulus (presented on a mini monitor positioned in front of M2’s face) in the non-social control condition (**Fig. 1a**, right). RDM stimulus was chosen as a non-social control since it has no behavioral meaning (i.e. no intrinsic value) to monkeys. On each experimental day, microstimulations were applied to one of the three prefrontal areas: OFC, dmPFC, or ACCg (**Fig. 1b–d**; **Fig. S1a**; Methods).

**Figure 1.**
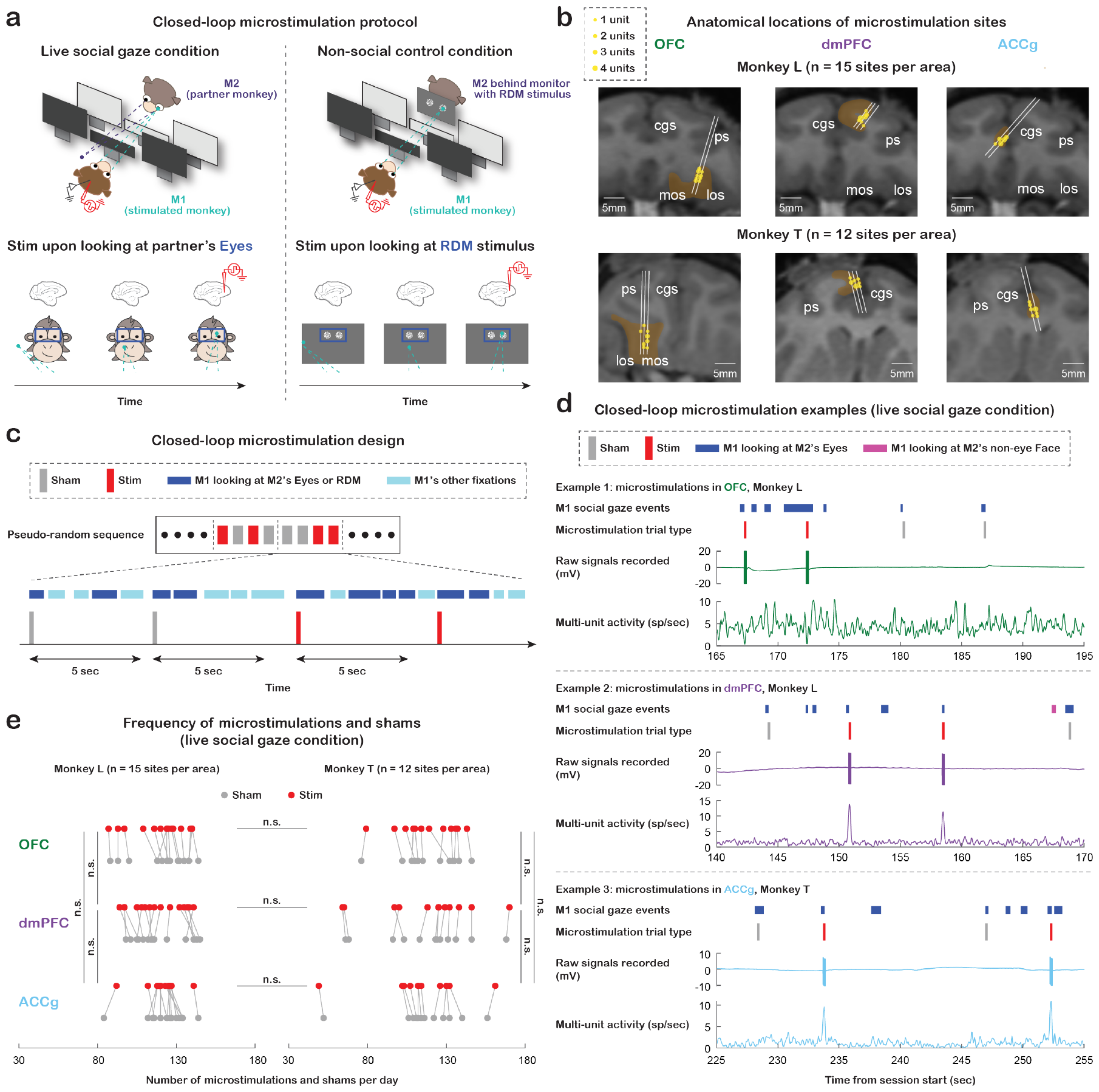
Experimental setup and microstimulation design. (**a**) Experimental paradigm for studying the functional role of the primate prefrontal cortex in naturalistic social gaze interaction. Left, live social gaze condition where each real-time microstimulation was selectively triggered by M1 fixating on M2’s *Eyes* for at least 30 msec with a probability of 50% (half microstimulation trials and half sham trials). Right, non-social gaze control condition where each real-time microstimulation was selectively triggered by M1 fixating for at least 30 msec on the *random dot motion (RDM) stimulus* (same location and size as *Eyes* ROI in the live social gaze condition) presented on a mini monitor placed in front of M2’s face. (**b**) Anatomical localizations of microstimulation sites in OFC, dmPFC, and ACCg from monkey L (n = 15 sites per area) and monkey T (n = 12 sites per area). (**c**) Diagram of the closed-loop microstimulation design. To avoid overstimulation of brain tissue, any two consecutive trials (including both microstimulations and shams) had to be at least 5 sec apart, and for every four trials, two microstimulations and two shams were randomly assigned. (**d**) Three examples of 30-sec experiment segments from the live social gaze condition. Each example, from top to bottom, shows M1’s *Eyes* (blue) and *non-eye Face* (pink) events (other fixations in space are not shown here), shams (gray) and microstimulations (red) triggered by looking at partner’s *Eyes*, raw signals recorded, and multi-unit activity. (**e**) Total number of microstimulations (red) and shams (gray) received per day in the live social gaze condition for monkey L (left) and monkey T (right). Data points connected with lines indicate measurements from the same day. The total number of microstimulations and shams per day was comparable across the three stimulated regions and comparable between the two animals (all p > 0.90). n.s., not significant, Wilcoxon rank sum, two-sided, FDR-corrected. Statistics for shams are not shown in the figure; none of the comparisons is significant.

The total number of microstimulations (and shams) received per day was comparable across the three stimulated regions and comparable between the two animals both in the social gaze and non-social control conditions (**Fig. 1e**, live social gaze condition, all p > 0.90, Wilcoxon rank sum, two-sided, FDR-corrected; **Fig. S1b**, non-social control condition, all p > 0.10). Further, we quantified spontaneously occurring gaze behaviors of the stimulated monkeys in the following regions of interest (ROIs): *Eyes* and *non-eye Face* (the rest of the face excluding the *Eyes* region) of the partner monkey in the live social gaze condition, and the *RDM stimulus* (same location and size as *Eyes* ROI) in the non-social gaze control condition. The total number of fixations on partner’s *Eyes* per day was significantly higher than fixations on *non-eye Face* for all three stimulated brain regions (**Fig. S1c**, top, all p < 10^-4^, Wilcoxon signed rank, two-sided, FDR-corrected), suggesting the significance of gaze directed to eyes that has been shown in previous studies in both humans and non-human primates ^8, 15, 16^. In addition, the total number of fixations on partner’s *Eyes* per day was comparable to fixations on the *RDM stimulus* for days involving the three stimulated regions (**Fig. S1c**, top, all p > 0.30), making it reasonable for us to compare the two conditions when examining microstimulation effect.

### Closed-loop microstimulations of OFC facilitate dynamic social attention in the spatial dimension

In our prior research, we elucidated a single-cell mechanism of social gaze monitoring in OFC, dmPFC, and ACCg. Notably, a significant proportion of neurons in these areas exhibited continuous and parametric tracking of where an individual is looking in space relative to another social agent or where the other agent is looking relative to oneself. This finding provides insight into a potential neural mechanism of social gaze monitoring involving these prefrontal regions ^8^. The current study investigated whether these prefrontal regions causally regulate such social gaze tracking.

To address this question, we first constructed a fixation density map for each trial considering all fixations during the analyzed post-gaze epoch (within 1.5 sec after the onset of a microstimulation or sham; Methods) in the visual space surrounding the *Eyes* and *whole Face* (union of *Eyes* and *non-eye Face*) of the partner monkey. Differences in such fixation density maps between microstimulation and sham trial types revealed a potential role of OFC in modulating momentary dynamic social attention. Specifically, microstimulations of OFC led to more clustered subsequent gaze fixations around the partner monkey (**Fig. 2a**; see **Fig. S2a** for the results from individual stimulated monkeys). To quantify this effect, for each microstimulation or sham, we calculated the average Euclidean distance between each of the stimulated monkey’s gaze fixations during the post-gaze epoch and the center of partner’s *Eyes* in the live social gaze condition (**Fig. 2b**, left, social gaze distance; Methods) or the center of *RDM stimulus* in the non-social gaze control condition (**Fig. 2b**, right, non-social gaze distance). We then compared the average of these gaze distances per day between microstimulation and sham trial types for each stimulated brain region.

**Figure 2.**
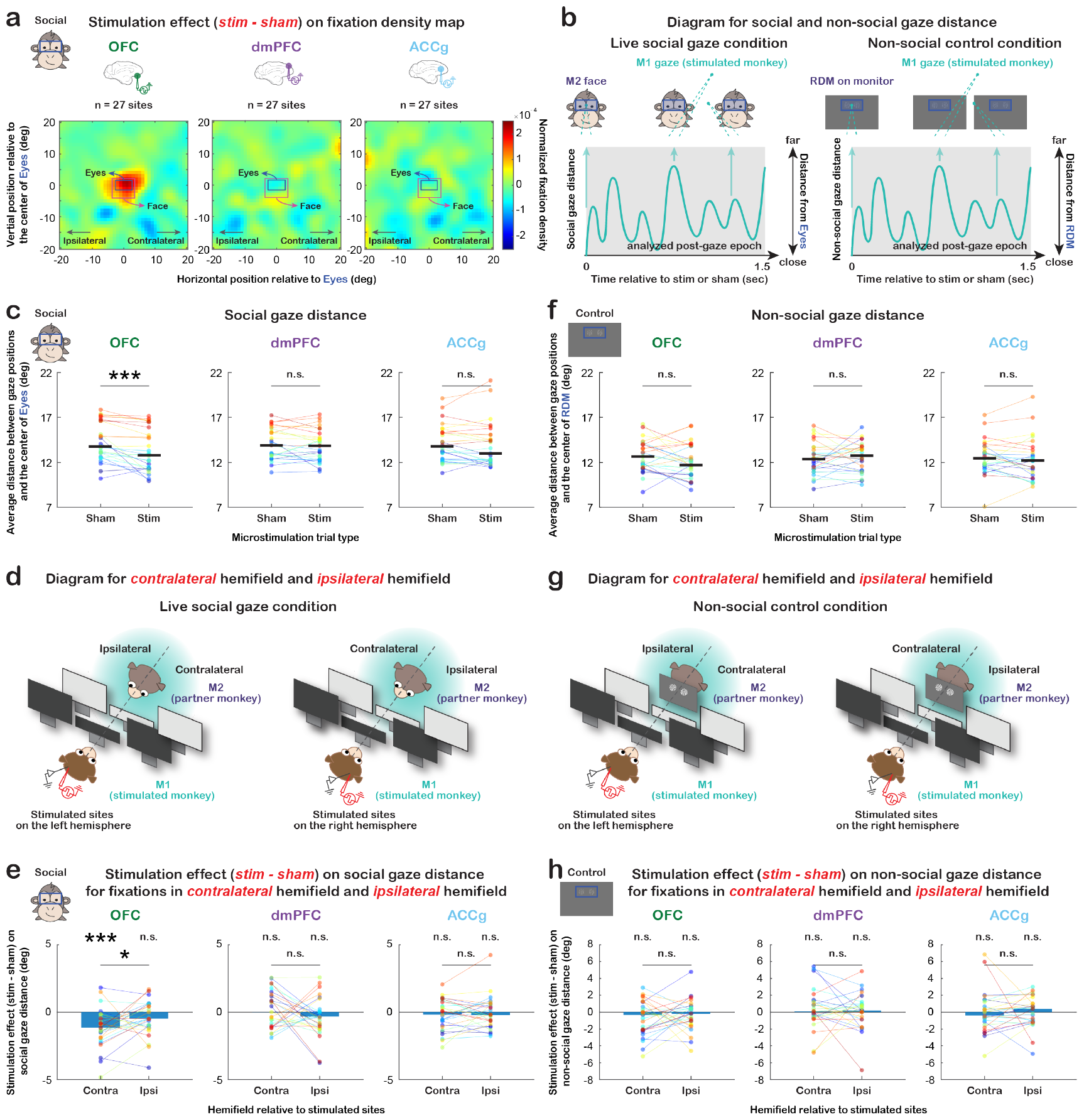
Microstimulation effects on dynamic social attention in the spatial dimension. (**a**) Microstimulation effect (difference between microstimulation and sham trial types) on the fixation density map of space surrounding partner monkey’s *Eyes* (blue rectangle) and *whole Face* (pink rectangle) for OFC, dmPFC, and ACCg (n = 27 sites per area). (**b**) Diagrams illustrating social and non-social gaze distances. For each microstimulation or sham, we calculated the average distance of all M1 fixations in space during the analyzed post-gaze epoch (within 1.5 sec after the onset of a microstimulation or sham) relative to M2’s *Eyes* in the live social gaze condition (social gaze distance, left) and relative to *RDM stimulus* in the non-social gaze control condition (non-social gaze distance, right). (**c**) Average social gaze distance per day (in visual degrees) for sham and microstimulation trial types separately for OFC, dmPFC, and ACCg. Data points in the same color connected with lines indicate measurements from the same day. Compared to shams, microstimulations of OFC significantly decreased social gaze distance during the post-gaze epoch (p < 0.001 for both monkeys combined; this effect was also present and significant in each monkey: p = 0.008 for monkey L and p = 0.002 for monkey T). *** p < 0.001, n.s., not significant, Wilcoxon signed rank, two-sided. (**d**) Diagrams illustrating the *contralateral* hemifield (opposite visual field of the stimulated brain hemisphere) and the *ipsilateral* hemifield (same visual field as the stimulated brain hemisphere) in the live social gaze condition. (**e**) Microstimulation effect on social gaze distance for fixations in the *contralateral* hemifield and *ipsilateral* hemifield separately for OFC, dmPFC, and ACCg. A negative value here (difference between microstimulation and sham trial types) indicates that microstimulations, compared to shams, resulted in more clustered subsequent gaze fixations around partner monkey’s *Eyes*. Data points in the same color connected with lines indicate measurements from the same day. The observed stimulation effect of OFC was more pronounced for gaze fixations in the contralateral visual field of the stimulated hemisphere (contralateral: p < 0.001 for both monkeys combined; p = 0.015 for monkey L and p = 0.012 for monkey T; ipsilateral: p = 0.068 for both combined; p = 0.208 for L and p = 0.233 for T; both hemifields combined: p < 0.001 for both combined; p = 0.008 for L and p = 0.002. for T). * p < 0.05, *** p < 0.001, n.s., not significant, Wilcoxon signed rank, two-sided. (**f–h**) Same format as (**c–e**) but for the non-social gaze control condition. n.s., not significant, Wilcoxon signed rank, two-sided.

As the fixation density maps show, microstimulations of OFC significantly decreased the average distance of one’s own gaze positions in space relative to partner’s *Eyes* during the post-gaze epoch, compared to shams (**Fig. 2c**, p < 0.001, Wilcoxon signed rank, two-sided). This suggests a facilitation of social attention in the spatial dimension by promoting gaze fixations around another social agent following OFC microstimulation. By contrast, we did not observe such stimulation effect on social gaze distance for dmPFC (**Fig. 2c**, p = 0.361) or ACCg (**Fig. 2c**, p = 0.374). Notably, the observed stimulation effect of OFC was more pronounced for gaze fixations in the contralateral visual field of the stimulated brain hemisphere (**Fig. 2d-e**; contralateral: p < 0.001; ipsilateral: p = 0.068; contralateral vs. ipsilateral: p = 0.026; Wilcoxon signed rank, two-sided). Again, no such effect was observed in either hemifield for dmPFC (**Fig. 2e**, contralateral: p = 0.501; ipsilateral: p = 0.149; contralateral vs. ipsilateral: p = 0.230) or ACCg (**Fig. 2e**, p = 0.442; p = 0.517; p = 0.564).

Crucially, these stimulation effects of OFC were exclusively observed in the live social gaze condition (i.e., microstimulations triggered by looking at partner’s *Eyes*) and not in the non-social gaze control condition using the *RDM stimulus* with no behavioral meaning to monkeys (**Fig. 2f**, OFC: p = 0.118; dmPFC: p = 0.719, ACCg: p = 0.302). The absence of stimulation effect for the *RDM stimulus* was also found when gaze fixation locations were split by hemifield for OFC (**Fig. 2g–h**; contralateral: p = 0.097; ipsilateral: p = 0.249; contralateral vs. ipsilateral: p = 0.442), dmPFC (p = 0.374; p = 0.517; p = 0.773) or ACCg (p = 0.532; p = 0.943; p = 0.171), supporting that the observed effects of OFC microstimulations in the spatial dimension were selective to social gaze interaction or when the stimulus had a behavioral meaning.

### Microstimulations of OFC also promote dynamic social attention in the temporal dimension

#### Inter-looking interval

In addition to the spatial dimension, the temporal aspect of social attention plays a crucial role in guiding social gaze interaction. Specifically, the time elapsed between individual instances of looking at another agent could serve as an index of social attention, with shorter durations between such gaze events indicating increased social attention. In this context, we sought to determine whether OFC microstimulations contributed to a reduction in the interval between social gaze events, in addition to the observed enhancement of social attention in the spatial dimension. Specifically, we examined the latency of M1 to look back at M2’s *whole Face* (i.e., the first *whole Face* event within 5 sec after the onset of a microstimulation or sham that was triggered by fixation to partner’s *Eyes* in the live social gaze condition, *inter-looking interval*; **Fig. 3a**; Methods). Microstimulations of OFC decreased this inter-looking interval (**Fig. 3b**, p = 0.035, Wilcoxon signed rank, two-sided). However, we did not observe such stimulation effect for dmPFC (**Fig. 3b**, p = 0.792) or ACCg (**Fig. 3b**, p = 0.291). Further, this reduction of inter-looking interval from OFC microstimulations was specific to social attention as no effect was observed in the nonsocial gaze condition (OFC: p = 0.773; dmPFC: p = 0.080; ACCg: p = 0.943). It is worth noting that this analysis had a relative low number of relevant gaze events compared to social gaze distance data from the spatial dimension analysis (i.e., the stimulated monkey did not look back at the partner’s *whole Face* during the examined time window on 41% of microstimulation and sham trials per day on average). Nevertheless, when we combined all trials for each stimulated region, we still observed similar results to the day-level analysis above (**Fig. S2b**, OFC: p = 0.010; dmPFC: p = 0.301; ACCg: p = 0.602; Wilcoxon rank sum, two-sided). Microstimulations of OFC therefore tended to lead monkeys to look back at another social agent faster, which may facilitate social gaze monitoring and dynamic social attention.

**Figure 3.**
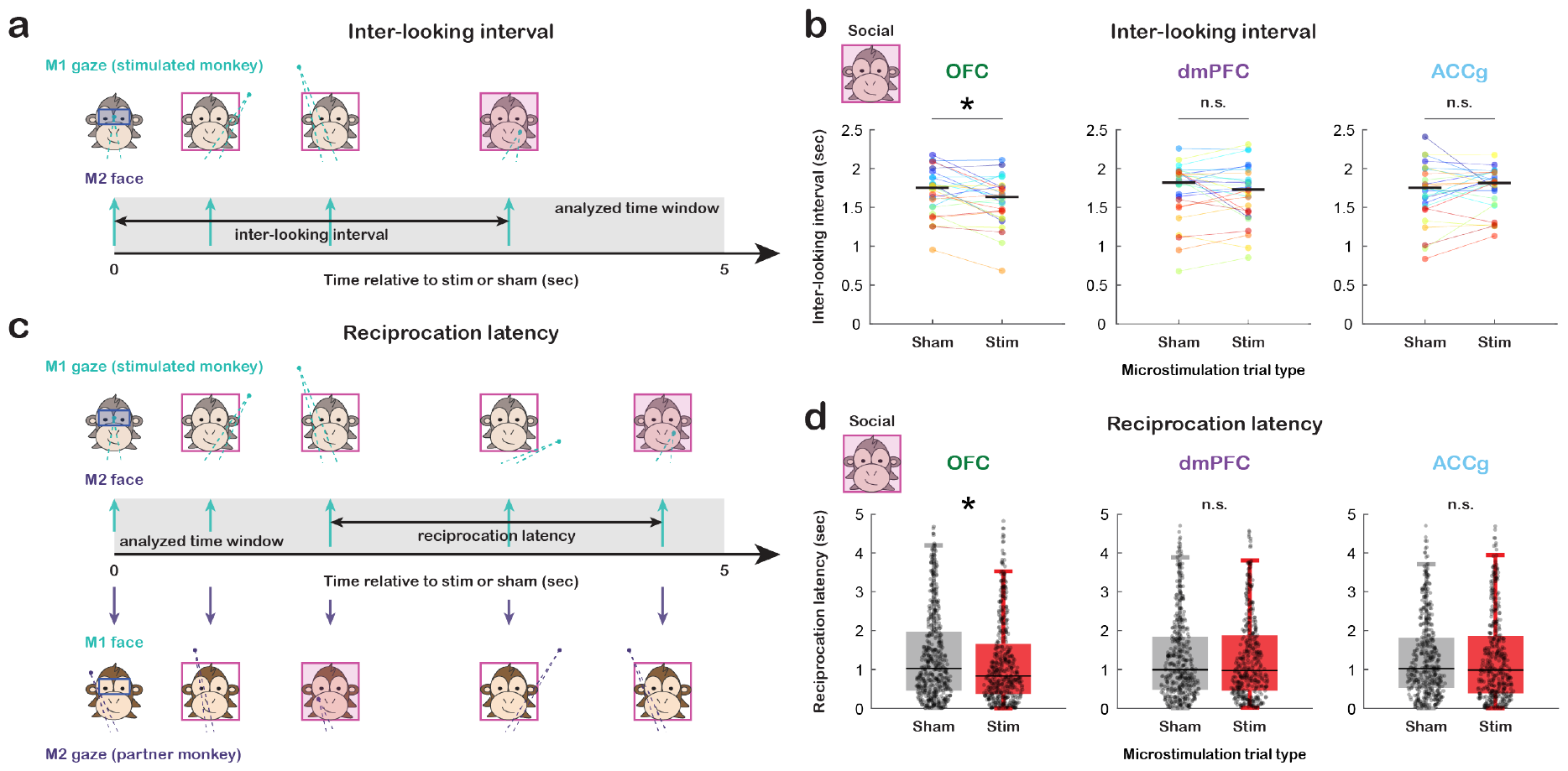
Microstimulation effects on dynamic social attention in the temporal dimension. (**a**) Diagram illustrating inter-looking interval, the latency of M1 to look back at M2’s *whole Face* during 5 sec after the onset of a microstimulation or sham. (**b**) Average inter-looking interval per day for sham and microstimulation trial types separately for OFC, dmPFC, and ACCg. Data points in the same color connected with lines indicate measurements from the same day. Microstimulations of OFC decreased inter-looking interval (p = 0.035 for both monkeys combined; p = 0.188 for monkey L and p = 0.092 for monkey T). * p < 0.05, n.s., not significant, Wilcoxon signed rank, twosided. (**c**) Diagram illustrating reciprocation latency, the latency of M1 to gaze back at M2’s *whole Face* after M2 looked at M1’s *whole Face* during 5 sec after the onset of a microstimulation or sham. (**d**) Distribution of reciprocation latency for sham (gray) and microstimulation (red) trial types separately for OFC, dmPFC, and ACCg. Trial-level data were collapsed across all days for each stimulated brain region. Microstimulations of OFC decreased reciprocation latency (p = 0.011 for both combined; p = 0.074 for L and p = 0.079 for T). * p < 0.05, n.s., not significant, Wilcoxon rank sum, two-sided.

#### Reciprocation latency

We next examined a more explicitly interactive aspect of social gaze dynamics. Specifically, we inspected the average latency of M1 to reciprocate gaze back at M2’s *whole Face* after M2 looked at M1’s *whole Face* within 5 sec after the onset of a microstimulation or sham that was triggered by fixation to partner’s *Eyes* (*reciprocation latency*; **Fig. 3c**; Methods). On the day level, microstimulations did not seem to greatly reduce such reciprocation latency (**Fig. S2c**, OFC: p = 0.130; dmPFC: p = 0.701; ACCg: p = 0.400; Wilcoxon signed rank, two-sided). However, this is likely due to a low number of relevant gaze events (i.e., there was no sequence of M2 looking at M1 and then M1 looking back at M2 during the examined time window on 86% of microstimulation and sham trials per day on average). When combining all trials for each stimulated region, we observed that microstimulations of OFC decreased reciprocation latency (**Fig. 3d**, p = 0.011, Wilcoxon rank sum, two-sided). We again did not observe such stimulation effect for dmPFC (**Fig. 3d**, p = 0.777) or ACCg (**Fig. 3d**, p = 0.368). Microstimulations of OFC therefore tended to lead monkeys to reciprocate another social agent’s gaze faster.

Thus, during spontaneous real-life social gaze interaction, closed-loop microstimulations of the OFC, following specific social gaze events, effectively enhanced momentary dynamic social attention in both spatial and temporal dimensions. In the spatial dimension, the subsequent gaze fixations were more clustered around another social agent, an effect more pronounced in the contralateral hemifield. In the temporal dimension, the inter-looking interval and reciprocation latency were shortened. Crucially, these effects were specific to social attention and were not observed for the *RDM stimulus*.

### Microstimulation effects of OFC are not driven by low-level properties of saccades

Importantly, the observed microstimulation effects of OFC were not driven by any change in the duration of the current gaze fixation to partner’s *Eyes* that itself triggered a microstimulation or sham (**Fig. S3a**, p = 0.302, Wilcoxon signed rank, two-sided), number of microsaccades (**Fig. S3b**, p = 0.456), number of macrosaccades (**Fig. S3c**, p = 0.055), macrosaccade kinematics indexed by saccade peak velocity over amplitude (**Fig. S3d**, p = 0.665, Wilcoxon signed rank, two-sided; **Fig. S3e**, p = 0.515, permutation test), or macrosaccade kinematics when considering saccade direction (**Fig. S3f**, ‘II’: macrosaccades from ipsilateral hemifield to ipsilateral hemifield; ‘IC’: macrosaccades from ipsilateral hemifield to contralateral hemifield; ‘CI’; ‘CC’; all p > 0.47, Wilcoxon signed rank, two-sided). Therefore, microstimulation effects of OFC were not associated with low-level changes in eye movements.

### Do microstimulations of OFC also lead to longer timescale modulation of social gaze exchanges?

The results reported above have shown that microstimulations of OFC enhance momentary dynamic social attention. Do these microstimulations also modulate social gaze exchanges on a longer timescale? To examine this, we analyzed inter-individual gaze dynamics between the stimulated monkey and the partner monkey. First, we applied a causal decomposition analysis ^17^ using moment-by-moment social gaze distance from each monkey (distance between one’s gaze positions and the center of the other monkey’s *Eyes*) during the post-gaze epoch (within 1.5 sec after the onset of a microstimulation or sham) and controlled for saccades (**Fig. 4a-b**; Methods). This allowed us to calculate a relative causal strength index that showed how much the gaze behaviors of one monkey in a pair was influenced by the gaze behaviors of the other monkey. To investigate stimulation effect on a longer timescale, we compared the first 45 stimulations (early epoch) to the next 45 stimulations (late epoch) from each day (Methods). While microstimulations of OFC enhanced momentary dynamic social attention as shown in the previous sections, they did not seem to impact gaze directionality indexed by the magnitude of relative causal strength for both time epochs combined (**Fig. S4a**, p = 0.239, Wilcoxon signed rank, two-sided) and for each epoch separately (**Fig. S4b**, all p > 0.16). Further, to examine whether inter-individual gaze dynamics were modulated by where oneself was looking in space, we correlated social gaze distance and relative causal strength (Methods) and found the slope of this fitted correlation comparable between early and late epochs for OFC (**Fig. S4c**, both hemifields combined: p = 0.757; **Fig. 4c**, contralateral: p = 0.882; **Fig. S4d**, ipsilateral: p = 0.098).

**Figure 4.**
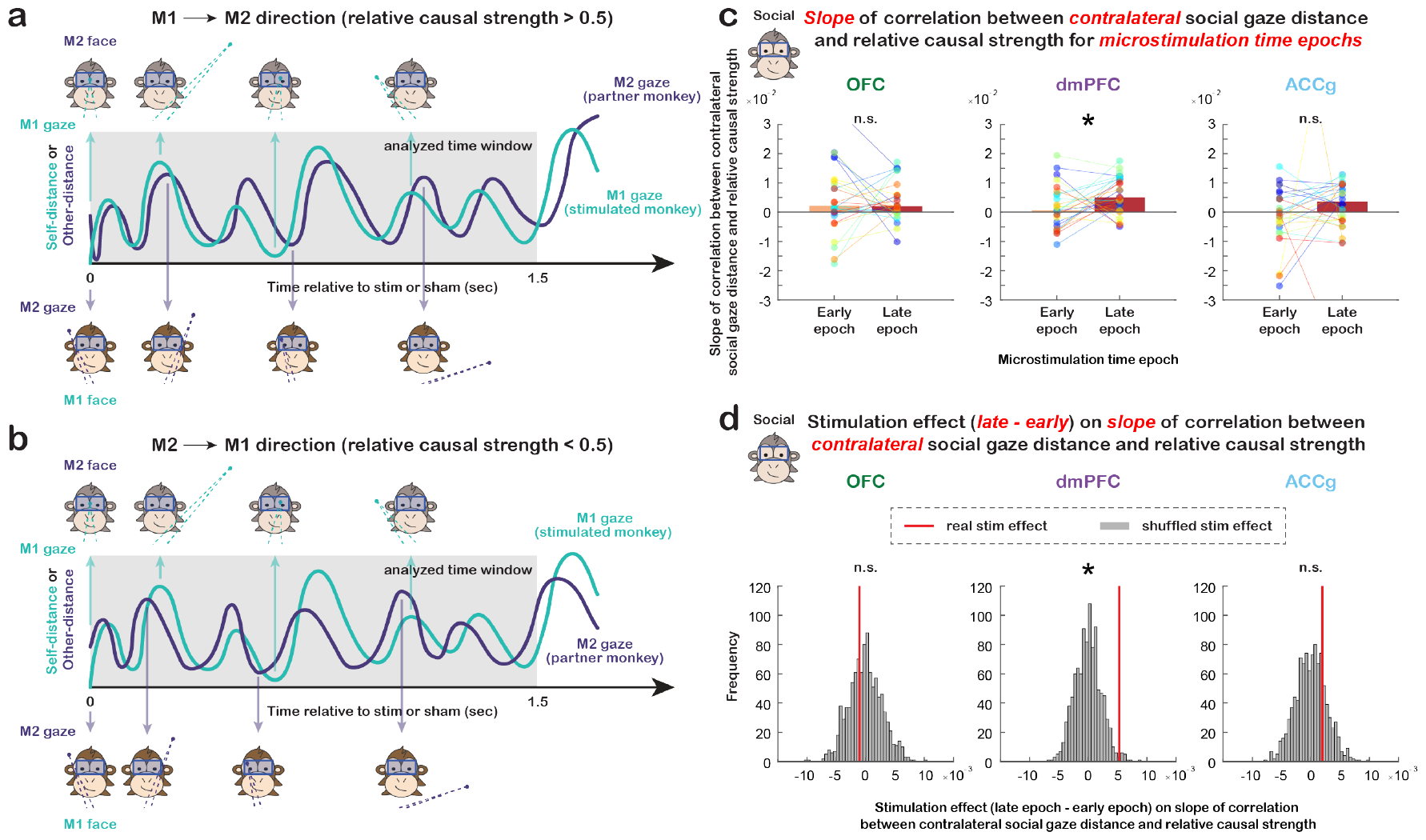
Longer timescale microstimulation effects on social gaze exchanges. (**a**) Diagram for M1 to M2 social gaze directionality when relative causal strength was greater than 0.5. (**b**) Diagram for M2 to M1 social gaze directionality when relative causal strength was less than 0.5. (**c**) Slope of correlation between social gaze distance in the contralateral hemifield and relative causal strength for microstimulations in the early epoch and late epoch separately for OFC, dmPFC, and ACCg. Data points in the same color connected with lines indicate measurements from the same day. For dmPFC, the slope of this fitted correlation was stronger for the late epoch than early epoch (p = 0.054 for both monkeys combined; p = 0.048 for monkey L and p = 0.625 for monkey T). * p ≈ 0.05, n.s., not significant, Wilcoxon signed rank, two-sided. (**d**) Microstimulation effect (difference between late and early time epochs) on the slope of examined correlation in (**c**). Red lines show the real median slope difference between late epoch and early epoch, whereas gray bars show the shuffled null distribution of slope difference medians (shuffling time epoch label 1,000 times for each day). The slope of this fitted correlation was stronger for the late epoch than early epoch for dmPFC when using gaze fixations in the contralateral hemifield (p = 0.015 for both combined; p = 0.038 for L and p = 0.068 for T). * p < 0.05, n.s., not significant, permutation test.

However, the slope of this fitted correlation for dmPFC was stronger for the late epoch than early epoch, specifically for gaze fixations in the contralateral hemifield (**Fig. 4d**; p = 0.015, permutation test; **Fig. S4e**, both hemifields combined: p = 0.131; **Fig. S4f**, ipsilateral: p = 0.455). These results suggested that microstimulations of dmPFC, but not OFC or ACCg, altered how social gaze exchanges were modulated by the location of the stimulated monkey’s gaze fixations in space on a longer timescale. Specifically, the slope of this examined correlation on average was positive in both early and late epochs for gaze fixations in both hemifields for dmPFC, indicating that as the stimulated monkey fixated closer around the partner monkey (smaller social gaze distance), his gaze behaviors were more likely to be led by the partner (lower relative causal strength). Intriguingly, as the number of dmPFC microstimulations accumulated within an experiment day, this effect became larger (greater slope for late epoch compared to early epoch). Microstimulations of dmPFC therefore altered how social gaze exchanges were modulated by the location of one’s gaze fixations on a relatively long timescale. These effects were also not driven by low-level properties of saccades (**Fig. S3**, all p > 0.16).

## Discussion

In primates, the gaze serves a critical function as they navigate through their social environment. Our previous electrophysiological work revealed that interactive social gaze variables are widely represented in the primate prefrontal-amygdala networks. In addition to the amygdala, a substantial proportion of neurons in OFC, dmPFC, and ACCg represent key signatures of social gaze interaction. Notably, spiking activity of many neurons in these prefrontal regions parametrically tracks one’s own gaze relative to another agent (‘social gaze distance’ also examined in the current paper) as well as other agent’s gaze relative to oneself ^8^. Here, we report that weak, realtime, closed-loop microstimulations of OFC modulate dynamic social attention. In the spatial dimension, these microstimulations resulted in clustered subsequent gaze fixations around another agent (reduced social gaze distance), an effect more pronounced for gaze fixations in the contralateral hemifield. In the temporal dimension, these microstimulations reduced the inter-looking interval for attending to another agent and the latency to reciprocate other’s directed gaze. These effects were found to be occurring on a relatively short timescale as OFC microstimulations did not change how long-term social gaze exchanges were modulated by the location of one’s own gaze fixations, unlike what we found with dmPFC microstimulations.

Widespread representations of social gaze variables in OFC, dmPFC, and ACCg neurons ^8^ are likely shaped by their common anatomical connectivity patterns with other brain regions in the social brain ^5^. The three prefrontal regions, albeit to different degrees, are bidirectionally connected to the amygdala ^18-20^, often referred to as the hub of social cognition ^21^ and implicated in both face and gaze processing ^8, 22-24^. Moreover, the orbitofrontal and medial prefrontal cortices, including the regions examined in this study, receive innervation from subregions in the inferior temporal cortex (IT) and the superior temporal sulcus (STS) ^25, 26^. These anatomical connections are likely to be functionally important for social gaze processing. The primate IT contains multiple face patches ^27, 28^ and the middle STS is believed to be a potential macaque homolog of the human temporal parietal junction, implicated in mentalizing in humans ^29^, based on both functional connectivity ^30^ and neural recoding ^31^. Face processing and mentalizing functions might be closely intertwined with the representations of social gaze variables. In this notion, it is possible that the interactive social gaze signals in OFC, dmPFC, and ACCg are subserving more abstract social cognitive functions that are functionally shared with social gaze processing.

During social gaze interaction, individuals constantly evaluate objects and other individuals in the environment and make momentary decisions to look toward or away from them. OFC neurons encode a wide range of outcome-related variables, such as expected value, choice value, reward prediction error, and choice and outcome history ^32-34^ that dynamically contribute to value and decision computations in OFC populations ^35-37^. These decision computations in OFC might facilitate the encoding of moment-to-moment value associated with other’s gaze and looking at other’s eyes for guiding adaptive behaviors. Indeed, value coding in OFC neurons is known to be modulated by gaze location. When monkeys freely viewed reward-predicting cues presented on a monitor, value signals in many OFC cells associated with the cues increased when monkeys fixated closer to the cues ^14^, suggesting a crucial role of OFC in both valuation and attention, two components foundational also to social gaze interaction. It has also been shown that in the OFC, weak microstimulations, similar to the ones used here, enhanced value computations during decision-making ^34^. Taken together, this might suggest a possible mechanism for the observed effects of OFC microstimulations. We hypothesize that weak, closed-loop microstimulations of OFC would increase the value signals associated with certain social gaze events and therefore enhance subsequent social attention.

It has been long theorized that looking at the face or the eyes of a conspecific has adaptive value ^38^. Indeed, value and social gaze variables have been shown to be representationally shared in the primate amygdala ^39^, which is strongly reciprocally connected to OFC ^20^. Our findings might reflect a synergistic effect of intrinsic value of social stimuli and microstimulation. Face and eyes are highly valued and readily capture attention. Weak microstimulations could further amplify the value signals in OFC ^34^ associated with looking at a social agent, thereby driving monkeys to fixate closer to and attend faster to the agent. Importantly, the effects of OFC microstimulations we observed were specific to the social context and not observed in the non-social control condition. This is likely because the *RDM stimulus* does not have any intrinsic or adaptive value in our experimental context, although it is visually salient and captures attention. We anticipate that OFC microstimulations would also enhance attention to certain non-social objects especially when they hold adaptive value, such as bananas, or learned cues that predict reward ^14^ or that guide gaze-following ^11^.

Studies that have causally manipulated activity in the primate brain have provided critical insights into brain functions. Microstimulations of the face patches in IT revealed their interconnectivity and distorted face perception^40^. Microstimulations of a gaze-following patch in the posterior STS impaired gaze-following behaviors when monkeys viewed images with different gaze directions ^11, 12^. In the decision-making literature, microstimulations of OFC were shown to bias choices ^34^. Further, closed-loop microstimulations of OFC delivered contingently upon theta frequency oscillation were shown to disrupt this synchronization and impair reward-guided learning ^41^. In the current work, closed-loop microstimulations of OFC delivered upon specific social gaze events enhanced dynamic social attention.

Intriguingly, we found that microstimulations of dmPFC altered how social gaze exchanges were modulated by the location of one’s gaze fixations on a longer timescale. The closer the stimulated monkey looked near the partner monkey, the more likely his gaze behaviors were led by the partner. Based on the hypothesized role of dmPFC in mentalizing and representing social information about self and other ^42-44^, this finding is consistent with the possibility that dmPFC microstimulations might have modulated the computations for understanding the intention of other’s gaze, which likely requires building an internal model of a social agent over multiple interactive bouts on a longer timescale. On the other hand, we observed neither short-term nor long-term microstimulation effect in ACCg. Given that social gaze signals are widely found in OFC, dmPFC, and ACCg ^8^, such dissociated functional consequences among the three areas from our closed-loop microstimulation protocol suggest potential differentiations on how neuromodulations affect different prefrontal populations. However, the current study cannot rule out if evoking behavioral changes in different neural tissues may require tailored stimulation protocols.

Future studies applying different stimulation protocols (non-closed-loop or closed-loop stimulations contingently upon a different gaze behavior of the stimulated monkey or specific gaze behavior of the social partner) might further reveal new insights into how the prefrontal cortex is involved in social gaze interaction. It is also critical to explore the stimulation parameter space as positive effects of different brain areas could depend on the choice of parameters ^34^. Moreover, in the current study, we used the same partner monkey who was familiar to the two stimulated monkeys. Social gaze dynamics have been shown to be influenced by dominance and familiarity ^15^. Further, the orbitofrontal and medial prefrontal networks are differentially connected to a specific region in the temporal pole ^45^ that processes personally familiar faces ^46^. Therefore, it would be informative to test if the observed effects of OFC microstimulations are modulated by social context.

During ongoing gaze exchanges, it is critical to dynamically increase or decrease attention to another social agent following specific social gaze events. Such behavioral contingency or adaptability is essential in guiding social interaction. Moreover, given the importance of social gaze in multitudes of social behaviors in primate species, social gaze representations in the brain may be tightly coupled to action or outcome related information about other social agents that are critical for observational learning and social decision-making ^42, 43, 47, 48^. Importantly, atypical visual attention and social gaze patterns are frequently associated with social disorders, such as autism spectrum disorder (ASD) ^49-51^. Our findings also have a therapeutic implication for using closed-loop microstimulation protocols – a ‘social brain interface’ – to modulate atypical social attention and social gaze behaviors in ASD. Stimulating OFC during an eye looking training session may help improve social attention, and stimulating dmPFC could potentially enhance responsiveness in social gaze exchanges on a longer timescale. Future investigations utilizing a noninvasive closed-loop stimulation protocol will help develop therapies to mitigate atypical social gaze behaviors.

## Acknowledgments

This work was supported by the National Institute of Mental Health (R01MH110750; R01MH120081; R01 MH128190).

## Author Contributions

S.W.C.C., S.F., and O.D.M. designed the study and wrote the paper. S.F., A.R.N., and O.D.M. performed the experiments. S.F., N.A.F., O.D.M., and S.W.C.C. analyzed the data.

## Methods

### Animals

Two adult male rhesus macaques (*Macaca mulatta*) were involved as stimulated monkeys (M1; monkeys L and T; both aged 10 years, weighing 15.7 kg and 14.1 kg, respectively). For each M1, unrelated monkey E (female, aged 10 years, weighing 10.9 kg) served as a partner monkey (M2). M2 was previously housed in the same colony room with M1s and other rhesus macaques and later moved to an adjacent colony room. The focus of the current study was to investigate the causal functions of OFC, dmPFC, and ACCg in dynamic social attention and did not include the necessary number of pairs to examine the effects of social relationships. Our previous published work using the identical paradigm has provided a comprehensive examination of the effects of social relationship on social gaze interaction from unique 8 dominance-related, 20 familiarity-related, and 20 sex-related perspectives ^15^. In this study, all animals were kept on a 12-hr light/dark cycle with unrestricted access to food, but controlled access to fluid during testing. All procedures were approved by the Yale Institutional Animal Care and Use Committee and in compliance with the National Institutes of Health Guide for the Care and Use of Laboratory Animals. No animals were excluded from our analyses.

### Experimental setup

On each day, M1 and M2 sat in primate chairs (Precision Engineering, Inc.) facing each other, 100 cm apart and the top of each monkey’s head 75 cm from the floor, with three monitors facing each monkey and the middle monitor 36 cm away from each monkey’s eyes (**Fig. 1a**). Two infrared eye-tracking cameras (EyeLink 1000, SR Research) simultaneously and continuously recorded the horizontal and vertical eye positions from both monkeys at 1,000 Hz. We conducted a two-step calibration procedure described in our previous work ^8^.

Each data collection day consisted of a total of alternating 10 *live social gaze* sessions and 5 *non-social control* sessions on average for monkey L (9-11 social and 4-8 control sessions across all days) and alternating 15 *live social gaze* sessions and 5 *non-social control* sessions on average for monkey T (14-15 social and 5 control sessions across all days). Each session lasted 300 sec. During *live social gaze* sessions, pairs of monkeys were allowed to freely interact with each other using gaze (**Fig. 1a**, left). During *non-social control* sessions, M1 was allowed to freely examine the space where a *random dot motion (RDM) stimulus* was presented on a mini monitor positioned on M2’s primate chair directly in front of M2’s face (**Fig. 1a**, right). At the beginning of each *live social gaze* session, the middle monitors were lowered down remotely so that the two monkeys could fully see each other (**Fig. 1a**, top). Before the beginning of each *non-social control* session, the mini monitor was positioned by an experimenter in front of M2’s face (**Fig. 1a**, right) and the middle monitors were lowered down remotely once the experimenter left the testing room. The mini monitor was 38 cm x 21 cm (W x H) at a resolution of 1024 pixel x 768 pixel. *RDM stimulus* was constructed using Variable Coherence Random Dot Motion MATLAB library (https://shadlenlab.columbia.edu/resources/VCRDM.html) and contained randomly moving dots within two circular apertures of 2.4 deg diameter each, with an inter-aperture horizontal distance of 1.6 deg equidistantly placed to the left and right of the center of M2’s *Eyes* ROI. *RDM stimulus* (white dots on a black background, with a density of 16.7 dots/deg^2^ per second) generated apparent motion either upward or downward with a 100% coherence with a fixed velocity 2 deg/sec. Motion direction remained consistent within a session. At the end of each session, the middle monitors were raised up remotely and blocked the stimulated monkey’s visual access to the partner monkey or the *RDM stimulus* during a 180-sec break.

### Surgery and anatomical localization

All animals received a surgically implanted headpost (Grey Matter Research) for restraining their head movement. A second surgery was performed on the two M1 animals to implant a custom chamber (Rogue Research Inc.) to permit recording and microstimulation in OFC (11 and 13), dmPFC (6 and 8), and ACCg (24a, 24b and 32) ^52^. Placement of the chambers was guided by both structural magnetic resonance imaging (MRI, 3T Siemens) scans and stereotaxic coordinates. See **Fig. 1b** for microstimulation sites on representative MR slices from both monkeys.

### Closed-loop microstimulation protocol

On each day before data collection, a guide tube was used to penetrate intact dura and to guide a microstimulation electrode (median impedance 50 kΩ) and a recording electrode (tungsten, FHC Inc), which were remotely lowered by using a motorized multi-electrode microdrive system (NaN Instruments) at the speed of 0.02 mm/sec. After the two electrodes reached target site, we ensured that we positioned the electrodes in the grey matter and waited 30 min for the tissue to settle for signal stability before starting experiment. The microstimulation site was usually positioned within 1mm from the recording electrode site on the chamber grid. Each closed-loop microstimulation (PlexStim system, Plexon Inc) was selectively triggered by M1 fixating within M2’s *Eyes* region for at least 30 msec in the live social gaze condition or M1 fixating within the *RDM stimulus* region (same location and size as *Eyes* ROI in the live social gaze condition) for at least 30 msec in the non-social control condition, with a probability of 50%. For every four trials, two microstimulations and two shams were randomly assigned. Parameters of each microstimulation (cathode-leading bipolar with a phase duration of 200 µsec and an interphase duration of 100 µsec) were 75 µA in amplitude, 100 Hz in frequency, and 200 msec in duration. Because the gaze-contingency of a microstimulation or sham was computed and implemented on the same microcontroller (Arduino), there was a negligible (< 1 msec) delay between registering a gaze fixation and initiating a microstimulation or sham. To avoid overstimulation of brain tissue, any two consecutive trials (including both microstimulations and shams) had to be at least 5 sec apart (**Fig. 1c-d**; **Fig. S1a**).

### Frequency of microstimulations and regions of interest for social and non-social gaze sessions

To quantify the frequency of microstimulations received by the stimulated monkeys, we calculated and compared the total number of microstimulations and shams per day across the three stimulated brain regions and two animals by using Wilcoxon rank sum test (**Fig. 1e**; **Fig. S1b**). On each experiment day, we identified the following regions of interest (ROIs): *Eyes* and *non-eye Face* (the rest of the face excluding the *Eyes* regions) in the live social gaze condition, and *RDM stimulus* in the non-social control condition (same location and size as *Eyes* ROI). In some analyses, we examined *whole Face* which is the union of *Eyes* and *non-eye Face* ROIs. Based on each day’s calibration, *whole Face* ROI was defined by the four corners of a monkey’s face and the *Eyes* ROI was defined by adding a padding of 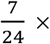 (width of the face − distance between the two eyes) to the center of each eye. Fixations were identified using EyeMMV toolbox in MATLAB ^53^. We detected fixations based on spatial and duration parameters, using t1 = 1.18 and t2 = 0.59 degrees of visual angle for the spatial tolerances and a minimum duration of 70 msec. As this fixation detection procedure does not incorporate velocity, we did not consider eye movement speed when identifying fixations. For each of the ROIs, we calculated the total number of gaze fixations and the average duration per fixation of the stimulated monkey for each day. Wilcoxon signed rank test was used to compare each variable between *Eyes* and *non-eye Face*, as well as *Eyes* and *RDM stimulus*. Wilcoxon rank sum test was used to compare each variable of each ROI between any pair of stimulated brain regions (**Fig. S1c**).

### Fixation density map

To construct a fixation density map, we examined the gaze positions of all M1’s fixations in space during the post-gaze epoch (within 1.5 sec after the onset of a microstimulation or sham) and assigned each fixation to a one-visual degree grid-square in a big spatial grid spanning 40 deg in both horizontal and vertical dimensions centered on partner’s *Eyes* ROI. Total number of fixations per day was calculated by summing across such fixations in each grid-square for microstimulation trials and sham trials separately and were z-scored. Difference in such fixation density between microstimulation and sham trial types was averaged across days for each stimulated brain region and plotted as heatmaps aligned to the center of partner’s *Eyes* (**Fig. 2a**, monkey T’s heatmaps were flipped horizontally as his chamber was implanted on the different hemisphere as monkey L; see **Fig. S2a** for individual stimulated animals). We focused on the post-gaze epoch as 1.5 sec was the common time window for both animals where we observed the reported stimulation effects (microstimulations of OFC in monkey L lasted for 3 sec after trial onset).

### Gaze distance analyses

In the live social gaze condition, for each trial, we examined all M1’s fixations during post-gaze epoch and calculated the Euclidean distance between each fixation and the center of partner’s *Eyes* ROI projected onto the same plane. Such distance was first averaged across all fixations after each trial and then averaged for each trial type for each day (**Fig. 2b**). Wilcoxon signed rank test was used to compare social gaze distance between microstimulation and sham trial types (**Fig. 2c**). Fixations were further categorized into those on the *contralateral* hemifield (opposite visual field of the stimulated brain hemisphere; **Fig. 2d**) and *ipsilateral* hemifield (same visual field as the stimulated brain hemisphere) for each M1 separately. The microstimulation effect on social gaze distance was examined for fixations in the *contralateral* and *ipsilateral* hemifield separately (**Fig. 2e**). Wilcoxon signed rank test was used to compare microstimulation effect on social gaze distance for each hemifield separately and between hemifields. The same analysis was applied for the non-social control condition (**Fig. 2f-h**) by calculating the Euclidean distance between M1’s fixations and the center of *RDM stimulus*.

### Gaze latency analyses

To examine dynamic social attention in the temporal dimension, we examined two measurements, *inter-looking interval* and *reciprocation latency*. Specifically, we focused on the *whole Face* ROI and combined events when only M1 fixated on the *whole Face* of M2 and the events when both monkeys’ gaze positions were within each other’s *whole Face*. First, we examined *inter-looking interval*, the latency for M1 to look back at M2’s *whole Face* within 5 sec after the onset of a microstimulation or sham (**Fig. 3a**). *Inter-looking interval* was calculated for each trial and averaged across all microstimulation trials and sham trials separately for each day (**Fig. 3b**). Wilcoxon signed rank test was used to compare such interval between microstimulation and sham trial types. Same analysis was applied for the non-social control condition by using the corresponding *whole Face* ROI. We also performed a trial-level analysis given that there was a relative low number of relevant gaze events by collapsing all microstimulation trials and all sham trials separately across all days for each stimulated brain region (**Fig. S2b**). Wilcoxon rank sum test was used to compare *inter-looking interval* between microstimulation and sham trial types on the trial level.

We next examined *reciprocation latency*, the latency for M1 to look back at M2’s *whole Face* after M2 looked at M1’s *whole Face* within 5 sec after the onset of a microstimulation or sham (**Fig. 3c**). *Reciprocation latency* was calculated for each trial and averaged across all microstimulation trials and sham trials separately for each day (**Fig. S2c**). Wilcoxon signed rank test was used to compare such latency between microstimulation and sham trial types. Again, due to the scarcity of relevant gaze events, we collapsed all microstimulation trials and all sham trials separately across all days for each stimulated brain region (**Fig. 3d**). Wilcoxon rank sum test was used to compare such latency between microstimulation and sham trial types. This analysis can only be applied in the live social gaze condition because there was no information about M2’s gaze in the non-social control condition given that M2’s visual access to M1 was blocked by the mini monitor placed in front of her.

### Saccade kinematics

To inspect if microstimulations resulted in any change in the current gaze event or saccade kinematics in the live social gaze condition, we first examined the average duration of current looking at partner’s *Eyes* that triggered a microstimulation or sham, as well as the total number of microsaccades and macrosaccades (**Fig. S3a-c**). These measurements were calculated for each day and Wilcoxon signed rank test was used to compare them between microstimulation and sham trial types for each stimulated brain region. We identified saccades using an unsupervised clustering method ^54^ that captured both canonical microsaccades (small deviations in position within an epoch in which the eye is mostly steady) and macrosaccades (a more explicit saccade to a new spatial location). We thus separated microsaccades and macrosaccades in the following way: for each detected saccade event, if the event occurred strictly within an interval that was separately identified as a fixation ^53^, we classified the event as a microsaccade; otherwise, we classified the event as a macrosaccade. We then examined macrosaccade kinematics. We looked at M1’s macrosaccades during the post-gaze epoch for each microstimulation or sham and calculated peak velocity (deg/sec) and amplitude (deg) for the first macrosaccade. For each day, we then fit a linear regression between peak velocity and amplitude of all such saccades for microstimulation trials and sham trials separately and calculated the slope difference between the two trial types (**Fig. S3d**). Wilcoxon signed rank test was used to compare such slope difference to zero. Within each day, we also created a shuffled null distribution of such slope differences by shuffling trial type label 1,000 times and compared the real median slope difference to the 1,000 medians of slope difference from the shuffled null distribution (**Fig. S3e**; permutation test). Furthermore, we categorized these macrosaccades into four groups depending on their direction (‘II’: saccades going from ipsilateral [I] hemifield to ipsilateral [I] hemifield; ‘IC’: saccades going from ipsilateral hemifield to contralateral [C] hemifield; also ‘CI’; ‘CC’). Wilcoxon signed rank test was used to test microstimulation effect on saccade kinematics against zero for each group separately.

### Inter-individual gaze dynamics analyses

In addition to M1’s gaze behaviors, we also examined how the two monkeys in a pair interacted with each other and their gaze directionality. By using moment-by-moment social gaze distance during the post-gaze epoch from each monkey (distance between one’s gaze positions and the center of the other monkey’s *Eyes*), we applied causal decomposition analysis ^17^ and calculated the average relative causal strength across all intrinsic mode functions (IMFs) for each trial. A relative causal strength value closer to 1 means stronger directionality from M1 (stimulated monkey) to M2 (partner monkey) and a value closer to 0 means stronger directionality from M2 to M1 (**Fig. 4a-b**). Because the causal decomposition analysis required continuous data, we smoothed gaze data to fill in the gaps between fixations and excluded a trial if more than 1 sec continuous eye tracking samples of either monkey were ‘NaN’ or the start and end points of either monkey’s smoothed portion were more than 20 visual degrees apart. Wilcoxon signed rank test was used to compare the relative causal strength between microstimulation and sham trial types (**Fig. S4a**). Again, this analysis can only be applied in the live social gaze condition as there was no information about M2’s gaze in the non-social control condition.

To further investigate microstimulation effect on a longer timescale, we divided microstimulations into early epoch (the first 45 stimulations) and late epoch (the next 45 stimulations) for each day. 76 out of 81 days had at least 90 stimulations. The 5 days excluded from further analysis were 1 day from monkey L with OFC microstimulations, 1 day from monkey T OFC, 2 days from monkey T dmPFC, and 1 day from monkey T ACCg. Wilcoxon signed rank test was used to compare the relative causal strength for different combinations of microstimulation trial types and time epochs (**Fig. S4b**). Then, for each day, we fitted a linear regression between social gaze distance and relative gaze causal strength. This analysis was conducted by using social gaze distance in both hemifields combined, contralateral hemifield, and ipsilateral hemifield. Wilcoxon signed rank test was used to compare the slope of this fitted line between late epoch and early epoch separately for OFC, dmPFC, and ACCg (**Fig. S4c**; **Fig. 4c**; **Fig. S4d**). Lastly, to make sure the observed results were robust, we calculated the real median slope difference between late epoch and early epoch for each stimulated region and compared it to the null distribution of slope difference medians by shuffling the temporal order of the 90 microstimulations for 1,000 times for each day (**Fig. S4e**; **Fig. 4d**; **Fig. S4f**; permutation test).

## Data availability

Behavioral and neural data presented in this paper will be available at the DANDI archive (changlabneuro).

## Code availability

Behavioral and neural data analysis codes central to this paper are available at https://github.com/changlabneuro/TBD.

## Supplementary Results

### Additional analyses and statistics on the spatial dimension of dynamic social attention

We focused on the post-gaze epoch (within 1.5 sec after the onset of a microstimulation or sham) because it was the common time window for both animals where we observed a significant decrease in social gaze distance following OFC microstimulations. In fact, this effect was present and lasted longer beyond 1.5 sec in one of the stimulated monkeys (within 2 sec after trial onset: p = 0.003 for both monkeys combined; p = 0.008 for monkey L and p = 0.204 for monkey T; within 3 sec: p = 0.004 for both combined; p = 0.007 for L and p = 0.204 for T).

In addition to looking at social gaze distance in a continuous manner, we also examined fixations in a binary fashion (a fixation within an ROI or not). Following OFC microstimulations, unlike more clustered subsequent gaze fixations around another social agent, we did not observe any change in the total number of fixations within partner’s *Eyes* (within 1.5 sec: p > 0.18 for both monkey L and monkey T; 2 sec: p > 0.24; 3 sec: p > 0.30) or *whole Face* (within 1.5 sec: p > 0.12; 2 sec: p > 0.20; 3 sec: p > 0.22), suggesting that the enhanced social attention from OFC microstimulations was driven by having spatially closer gaze fixations around another social agent but not necessarily increased the number of fixations within the social agent’s eyes or face regions. However, this conclusion might be limited to the closed-loop microstimulation paradigm and to our specific stimulation parameters.

### Additional analyses and statistics on low-level properties of saccades

The observed effects of OFC and dmPFC microstimulations were not driven by any change in the duration of current looking to partner’s *Eyes* that triggered a microstimulation or sham (**Fig. S3a**, OFC: p = 0.302 for both monkeys combined; p = 0.679 for monkey L and p = 0.339 for monkey T; dmPFC: p = 0.269 for both combined; p = 0.107 for L and p = 0.970 for T; Wilcoxon signed rank, two-sided), number of microsaccades (**Fig. S3b**, OFC: p = 0.456 for both combined; p = 0.978 for L and p = 0.233 for T; dmPFC: p = 0.581 for both combined; p = 0.303 for L and p = 0.569 for T), number of macrosaccades (**Fig. S3c**, OFC: p = 0.055 for both combined; p = 0.188 for L and p = 0.110 for T; dmPFC: p = 0.230 for both combined; p = 0.978 for L and p = 0.003 for T), macrosaccade kinematics indexed by saccade peak velocity over amplitude (**Fig. S3d**, OFC: p = 0.665 for both combined; p = 0.762 for L and p = 0.424 for T; dmPFC: p = 0.904 for both combined; p = 0.639 for L and p = 0.569 for T; Wilcoxon signed rank, two-sided; **Fig. S3e**, OFC: p = 0.515 for both combined; p = 0.507 for L and p = 0.178 for T; dmPFC: p = 0.164 for both combined; p = 0.240 for L and p = 0.509 for T; permutation test), or macrosaccade kinematics when considering saccade direction (**Fig. S3f**, II: macrosaccades from ipsilateral hemifield to ipsilateral hemifield; IC: macrosaccades from ipsilateral hemifield to contralateral hemifield; CI; CC; OFC: all p > 0.47 for both combined; all p > 0.59 for L and all p > 0.12 for T; dmPFC: p > 0.31 for both combined; p > 0.30 for L and p > 0.26 for T; Wilcoxon signed rank, two-sided).

## Supplementary Figures and Legends

**Figure S1.**
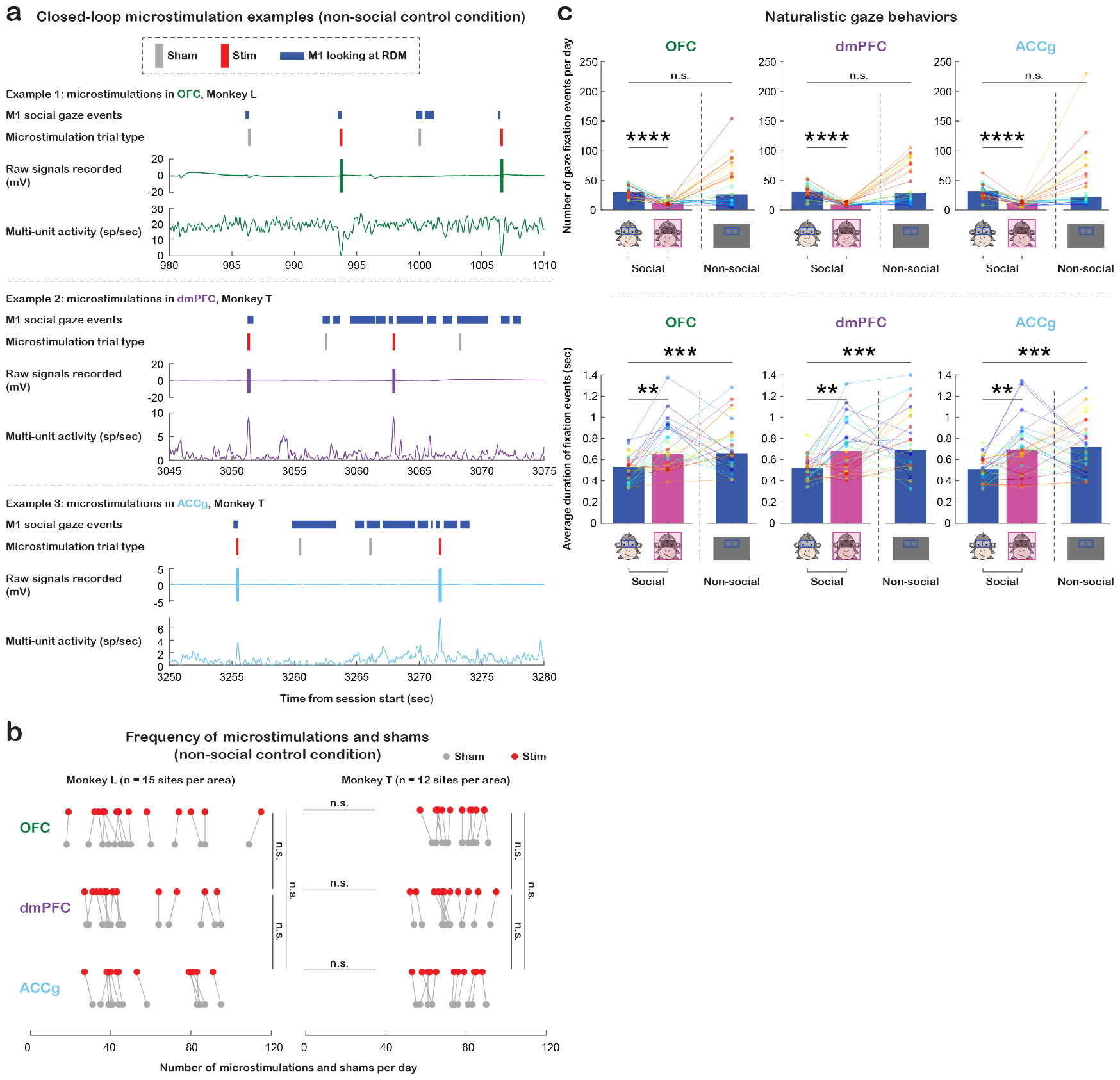
Additional analyses for the non-social control condition and naturalistic gaze behaviors. (**a**) Three examples of 30-sec experiment segments from the non-social control condition. Same format as **Fig. 1d**. Each example, from top to bottom, shows M1’s fixations on *RDM stimulus* (blue; other fixations in space are not shown here), realtime shams (gray) and microstimulations (red) triggered by looking at *RDM stimulus*, raw signals recorded, and multiunit activity. (**b**) Total number of microstimulations (red) and shams (gray) received per day in the non-social control condition for monkey L (left) and monkey T (right). Data points connected with lines indicate measurements from the same day. The total number of microstimulations and shams per day was comparable across the three stimulated regions and comparable between the two animals (all p > 0.10). n.s., not significant, Wilcoxon rank sum, two-sided, FDR-corrected. Statistics for shams are not shown in the figure; none of the comparisons is significant. (**c**) Naturalistic gaze behaviors summarized as the total number (top) and average duration per fixation (bottom) within partner monkey’s *Eyes* and *non-eye Face* in the live social gaze condition, as well as fixations to the *RDM stimulus* in the non-social control condition. Data points in the same color connected with lines indicate measurements from the same day. ** p < 0.01, *** p < 0.001, **** p < 0.0001, n.s., not significant, Wilcoxon signed rank, two-sided, FDR-corrected.

**Figure S2.**
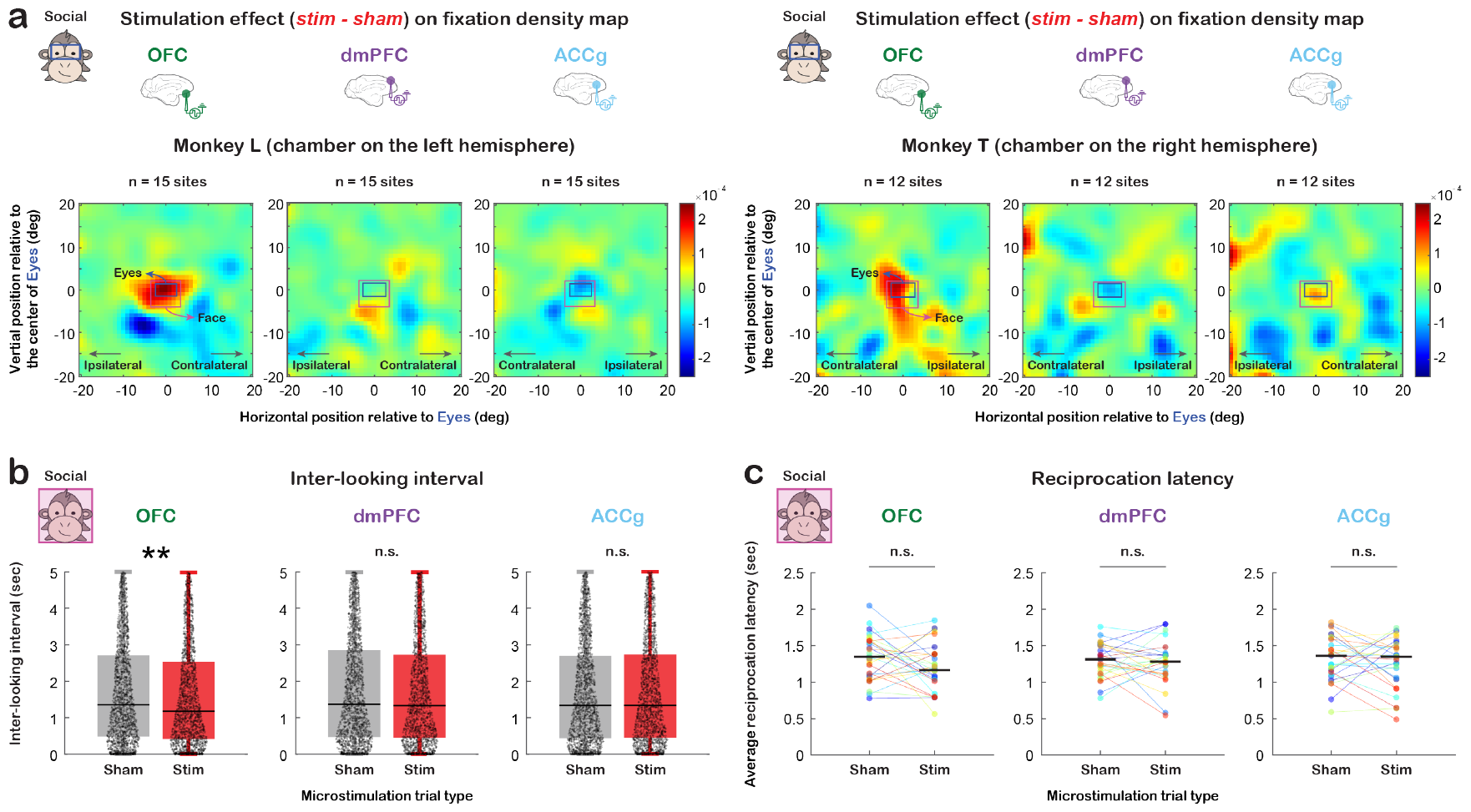
Additional analyses for dynamic social attention in the spatial and temporal dimensions. (**a**) Microstimulation effect (difference between microstimulation and sham trial types) shown on the fixation density map of space surrounding partner monkey’s *Eyes* (blue rectangle) and *whole Face* (pink rectangle) for OFC, dmPFC, and ACCg for monkey L (n = 15 sites per area, left) and monkey T (n = 12 sites per area, right) separately. Same format as **Fig. 2a**. (**b**) Distribution of inter-looking interval for sham (gray) and microstimulation (red) trial types separately for OFC, dmPFC, and ACCg. Trial-level data were collapsed across all days for each stimulated brain region. Microstimulations of OFC decreased inter-looking interval (p = 0.010 for both monkeys combined; p = 0.026 for monkey L and p = 0.143 for monkey T). ** p < 0.01, n.s., not significant, Wilcoxon rank sum, two-sided. (**c**) Average reciprocation latency per day for sham and microstimulation trial types separately for OFC, dmPFC, and ACCg. Data points in the same color connected with lines indicate measurements from the same day. On the day level, microstimulations did not seem to greatly reduce reciprocation latency (OFC: p = 0.130; dmPFC: p = 0.701; ACCg: p = 0.400). n.s., not significant, Wilcoxon signed rank, two-sided.

**Figure S3.**
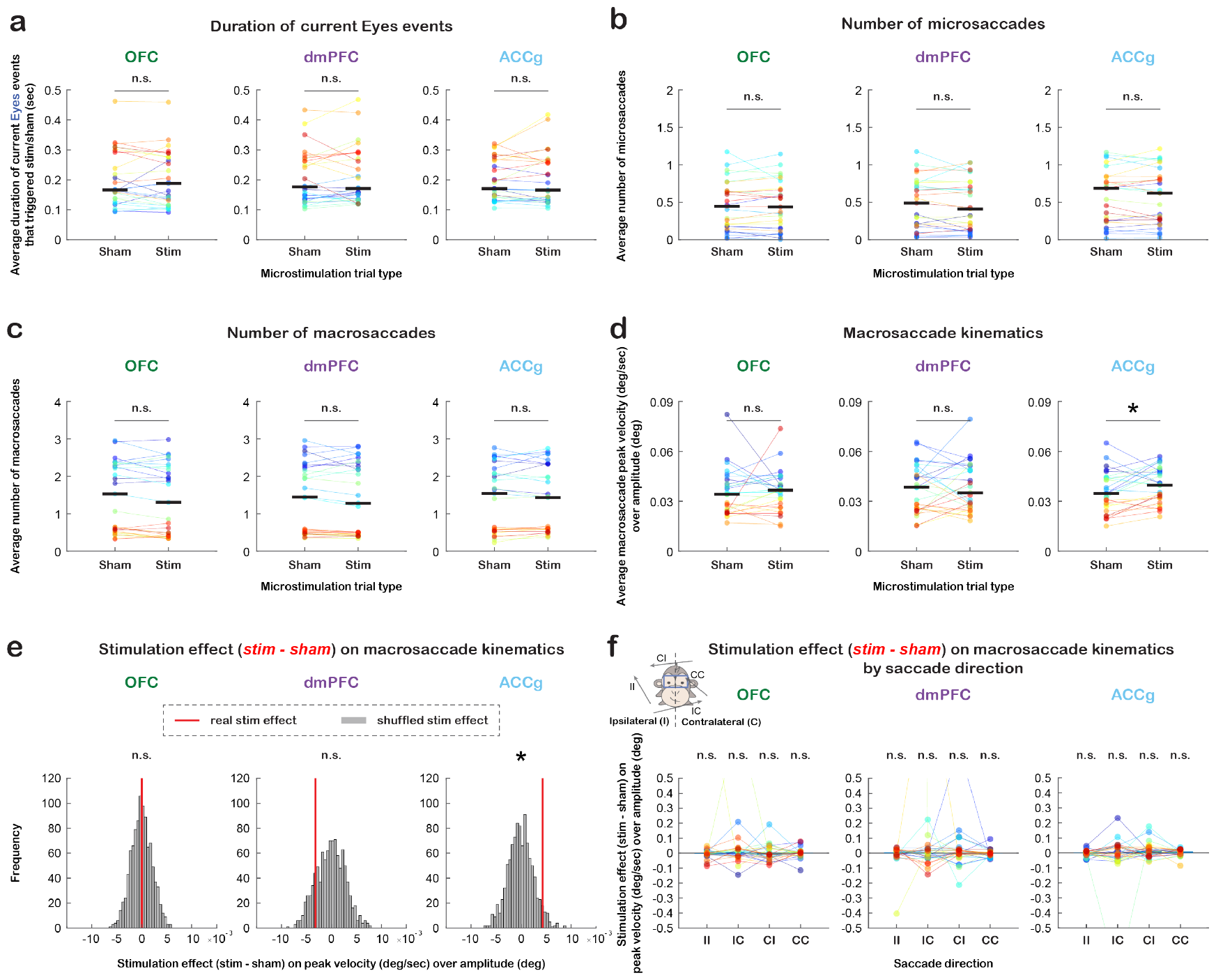
Control analyses on current gaze events and saccades. (**a**) Average duration per day of current *Eyes* events that triggered a microstimulation or sham, for sham and microstimulation trial types separately for OFC, dmPFC, and ACCg. Data points in the same color connected with lines indicate measurements from the same day. Please see Supplementary Results for more statistics. n.s., not significant, Wilcoxon signed rank, two-sided. (**b-c**) Average number of microsaccades (**b**) and macrosaccades (**c**) per day during post-gaze epoch for the two trial types separately for OFC, dmPFC, and ACCg. n.s., not significant, Wilcoxon signed rank, two-sided. (**d**) Average macrosaccade kinematics per day indexed by saccade peak velocity over amplitude for the two trial types separately for OFC, dmPFC, and ACCg. * p < 0.05, n.s., not significant, Wilcoxon signed rank, two-sided. (**e**) Microstimulation effect (difference between microstimulation and sham trial types) on macrosaccade kinematics. Red lines show the real median stimulation effect, whereas gray bars show the shuffled null distribution (shuffling microstimulation trial type label 1,000 times for each day). * p < 0.05, permutation test. (**f**) Microstimulation effect on macrosaccade kinematics by saccade direction separately for OFC, dmPFC, and ACCg. II: macrosaccades from ipsilateral hemifield to ipsilateral hemifield; IC: macrosaccades from ipsilateral hemifield to contralateral hemifield; CI; CC. n.s., not significant, Wilcoxon signed rank, two-sided.

**Figure S4.**
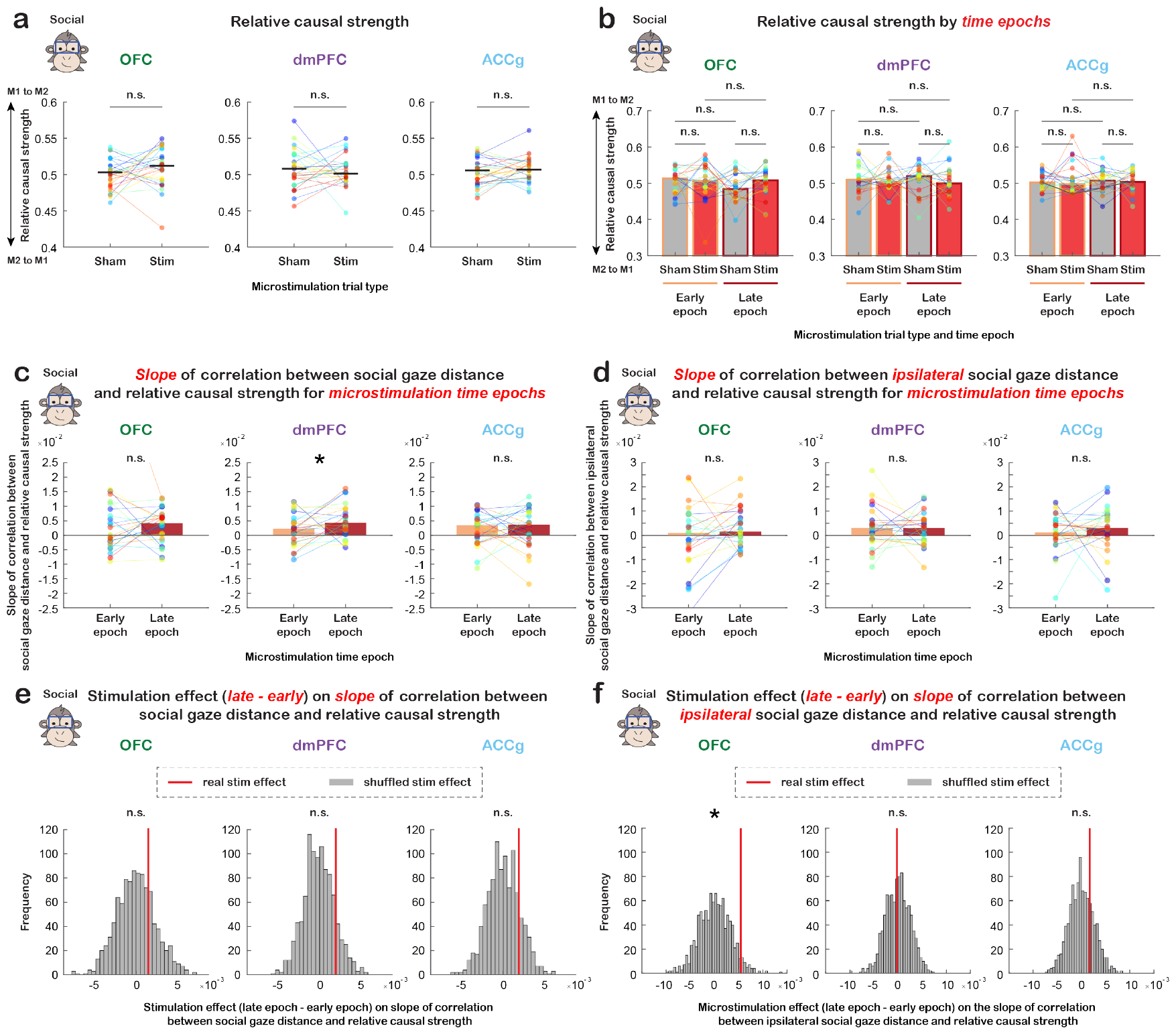
Additional analyses on longer timescale social gaze exchanges. (**a**) Average relative causal strength per day for sham and microstimulation trial types separately for OFC, dmPFC, and ACCg. Data points in the same color connected with lines indicate measurements from the same day. We did not observe stimulation effect on the magnitudes of relative causal strength (OFC: p = 0.239; dmPFC: p = 0.962; ACCg: p = 0.361; Wilcoxon signed rank, two-sided). n.s., not significant, Wilcoxon signed rank, two-sided. (**b**) Average relative causal strength per day for different microstimulation trial types and time epochs (orange: early epoch; red: late epoch) separately for OFC, dmPFC, and ACCg. Data points in the same color connected with lines indicate measurements from the same day. We did not observe effect of microstimulation trial type or time epoch on relative casual strength (OFC: all p > 0.16; dmPFC: all p > 0.67; ACCg: all p > 0.46). n.s., not significant, Wilcoxon signed rank, two-sided. (**c**) Slope of correlation between social gaze distance and relative causal strength for early epoch and late epoch separately for OFC, dmPFC, and ACCg. The slope of this fitted correlation was stronger for the late epoch than early epoch for dmPFC, but not for the other two regions (dmPFC: p = 0.037; OFC: p = 0.757; ACCg: p = 0.770). * p < 0.05, n.s., not significant, Wilcoxon signed rank, two-sided. (**d**) Same format as (**c**) but when using social gaze distance in the ipsilateral hemifield. The slope of this fitted correlation was comparable between the two time epochs (OFC: p = 0.098; dmPFC: p = 0.757; ACCg: p = 0.381). n.s., not significant, Wilcoxon signed rank, two-sided. (**e**) Microstimulation effect (difference between late epoch and early epoch) on the slope of examined correlation in (**c**). Red lines show the real median slope difference between late epoch and early epoch, whereas gray bars show the shuffled null distribution of slope difference medians (shuffling time epoch label 1,000 times for each day). n.s., not significant, permutation test. (**f**) Same format as (**e**) but when using social gaze distance in the ipsilateral hemifield. * p < 0.05, n.s., not significant, permutation test.

